# Features of peptide fragmentation spectra in single cell proteomics

**DOI:** 10.1101/2021.08.17.456675

**Authors:** Hannah Boekweg, Daisha Van Der Watt, Thy Truong, Amanda J Guise, Edward D Plowey, Ryan T Kelly, Samuel H Payne

## Abstract

The goal of proteomics is to identify and quantify the complete set of proteins in a biological sample. Single cell proteomics specializes in identification and quantitation of proteins for individual cells, often used to elucidate cellular heterogeneity. The significant reduction in ions introduced into the mass spectrometer for single cell samples could impact the features of MS2 fragmentation spectra. As all peptide identification software tools have been developed on spectra from bulk samples and the associated ion rich spectra, the potential for spectral features to change is of great interest. We characterize the differences between single cell spectra and bulk spectra by examining three fundamental spectral features that are likely to affect peptide identification performance. All features show significant changes in single cell spectra, including loss of annotated fragment ions, blurring signal and background peaks due to diminishing ion intensity and distinct fragmentation pattern compared to bulk spectra. As each of these features is a foundational part of peptide identification algorithms, it is critical to adjust algorithms to compensate for these losses.

## Introduction

Life is the result of dynamic interactions that occur within and among individual cells; the state of each cell and its ability to respond to environmental signals is determined by protein abundances. Therefore a primary goal of proteomics is to characterize the functional state of cells through the identification and quantification of proteins. Until recently, researchers have been limited to bulk proteomic analyses, in which a sample introduced to the mass spectrometer was derived from thousands or millions of cells. While this provides deep protein coverage, it obscures important heterogeneity that exists between individual cells. The only way to detect cellular heterogeneity is by profiling single cells^1–3^. Single cell proteomics (SCP) is rapidly advancing; it is now possible to quantify > 1000 proteins from a single cell^4–6^.

SCP presents unique technical challenges. For example, limited sample input leads to fewer ions accumulated in the mass spectrometer and lower signal intensity. Recent efforts to improve data quality for extremely low signal have focused on sample preparation^5,7^, chromatography and data acquisition^8,9^, and instrumentation^4,10^. Despite the excitement for SCP and the advances in data generation techniques, a relatively unexplored area is data analysis - specifically the fundamental algorithms for peptide identification.

Tandem mass spectrometry identification algorithms for proteomics match a peptide sequence to an observed spectrum and identify the most likely candidate peptide. Early peptide identification algorithms like SEQUEST^11^ relied on matching peaks of the observed spectra to the theoretical spectrum of fragment ions. The next generation of algorithms probabilistically score individual peaks based on expected intensities and isotope profiles. These techniques originated in *de novo* algorithms^12–14^, but are also used in database search tools^15^. A further improvement in spectral annotation comes from incorporating a rich set of spectral features for semi-supervised machine learning^16^. Algorithms for annotating mass spectra are continually evolving and some software tools have explicitly generated multiple scoring functions for improved performance with different fragmentation methods or instrument types^17^. The past 20-30 years of development in scoring functions have significantly improved peptide identification, but it’s important to note that they were all developed using data from bulk samples. Due to the recent emergence of single cell proteomics, no algorithm has specifically been developed for single cell data.

Single cell spectra potentially have different characteristics which might affect the success of software tools and scoring functions in identifying the best peptide/spectrum match (PSM). The following features are pillars of current algorithms: 1) the number of fragment ion peaks matched between the observed and theoretical spectrum, 2) peak intensity of fragment ions is expected to be much greater than background noise, and 3) repeated spectra of the same peptide will look similar. These features, and their derived metrics, are used to distinguish correct and incorrect PSMs. However, it is unknown how discriminating these features are in single cell data; understanding the data’s structure is essential for accurate software tools^12^. We explore these three features in single cell proteomics data and characterize how these spectra differ from bulk spectra. We discover fundamental differences that potentially impact the success of all peptide identification tools, specifically we find a dramatic loss of fragment ions, a compression of signal intensity and the blurring of signal and noise.

## Results

### Spectral features

Peptide identification algorithms rely on the features of fragmentation spectra, which may vary based on fragmentation method, instrument type or other experimental conditions such as ultra-low ion counts. To identify how spectral features change in single cell proteomics, we compared peptide/spectrum matches (PSMs) from three different HeLa datasets: a traditional bulk sample^18^, 2ng sample loading (∼10 cell equivalent), and 0.2ng sample loading (∼1 cell equivalent). Specifically, we characterize the number of annotated peaks, and intensity range of annotated and unannotated peaks. All spectra were searched using MetaMorpheus (see Methods) and the output was binned into two groups: high quality PSMs with q-value < 0.01 and mediocre PSMs with q-value between 1% and 20%. Throughout the manuscript these are referred to as ‘quality PSMs’ and ‘mediocre PSMs’.

To help establish an intuition for the spectral features discussed within this manuscript, Figure 1 shows three annotated spectra for the same peptide with features and associated metrics (see Methods). The first feature is the number of annotated *y* ion peaks. The spectrum from bulk proteomics (Fig 1A) has 10 annotated *y* ions; the 2ng quality spectrum (Fig 1B) has 8 annotated *y* ions; the 2ng mediocre spectrum (Fig 1C) has 5 annotated *y* ions. The second feature describes the range of intensity in a spectrum, a ratio of the average intensity of top 3 annotated peaks to the bottom 3 annotated peaks. The ratio for the bulk PSM, quality 2ng PSM and mediocre 2ng PSM in Figure 1 is 27, 5 and 2 respectively. A smaller ratio, like is found in the 2ng PSMs, demonstrates a compression of the intensity range within a spectrum. We also calculate a similar metric - the ratio between the average intensity of the top 3 annotated peaks and the median of all non-annotated peaks. MS2 spectra typically have many low intensity peaks which are not annotated and are considered background or noise. PSMs which have a low ratio for this metric are those for which annotated peaks and non-annotated peaks generally have a similar intensity (see Fig 1C).

**Figure 1.**
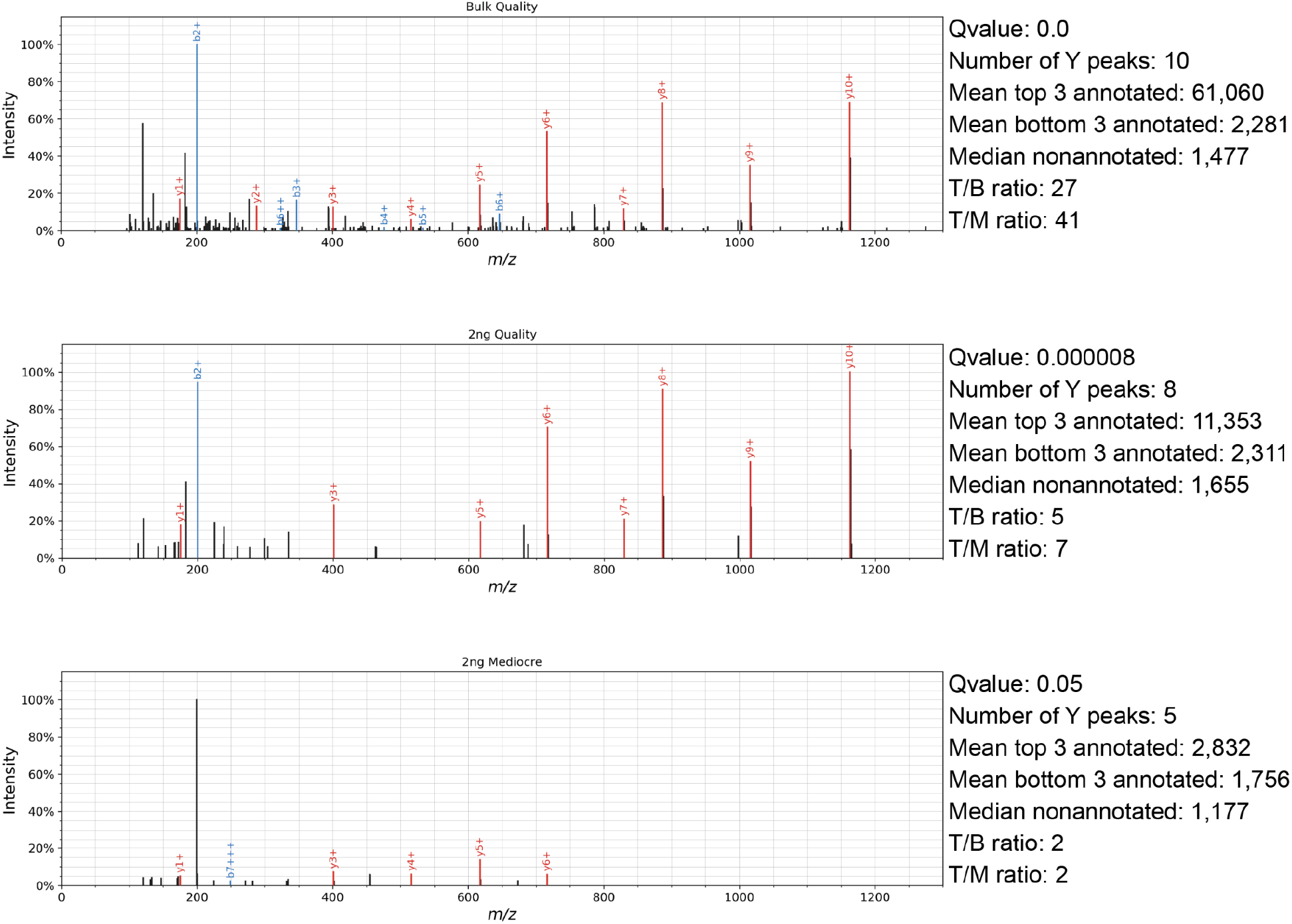
Data quality metrics for annotated spectra from bulk and single cell samples. All panels are spectra for the same peptide, AQFEGIVTDLIR. Each panel contains the annotated b/y peaks. Adjacent to the spectrum images are metrics which describe the quality of the peptide/spectrum match (PSM). The top panel is from bulk data; the middle panel is a high quality PSM from 2 ng data; the bottom panel is a mediocre PSM from 2 ng data. Note that most metrics are weaker for single cell PSMs. i.e. fewer annotated Y peaks and smaller intensity ratios.

### Peak loss in single cell data

Although algorithm implementation details differ for various peptide identification software tools, an algorithm will report a less confident score for PSMs that contain fewer annotated peaks. Therefore, it is essential to evaluate whether the number of annotated peaks in single cell spectra is similar to those in bulk proteomics data for which all algorithms have been developed. For peptides which had confident identifications in a bulk proteomics dataset, we compared the number of *y* ion peaks present in spectra from bulk proteomics, small input (2ng), and single cell proteomics (0.2 ng). Specifically, we calculated how many peaks are lost relative to PSMs from bulk samples (see Methods). As shown in Figure 2, confidently annotated spectra generally lose only a few annotated y ions. For PSMs with < 10 annotated y ions, this loss is generally 0-2 peaks. Larger peptides, whose spectra have more *y* ions tend to lose a few more peaks. For PSMs with a mediocre q-value, the number of annotated peaks is typically half as many as quality bulk data. For example, when looking at spectra that have an 10 annotated *y* ion peaks, quality single cell PSM loses ∼1 peak, and mediocre single cell PSM loses about 5. The results for 0.2 ng data were similar but with more pronounced peak loss (Supplemental Figure 1).

**Figure 2.**
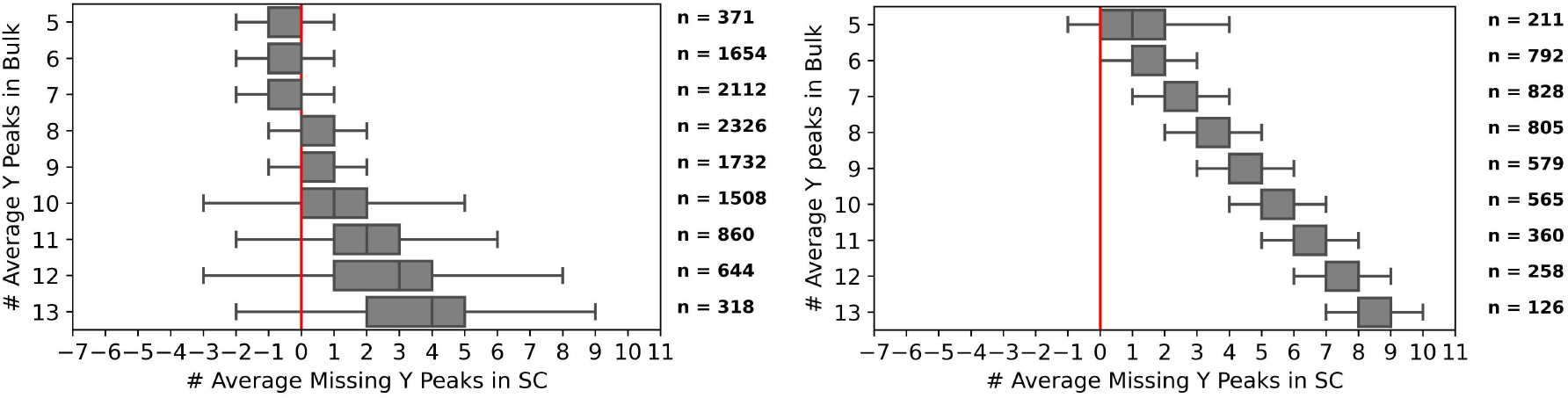
*Y* ion loss in single cell spectra. Plots show the number of peaks lost in single cell PSMs compared to the same peptide from bulk data. The 0 on the x axis (emphasized with a red line) indicates that the number of peaks is the same. Positive numbers indicate that there are missing peaks in single cell spectra compared to bulk spectra. A) peaks lost in quality single cell PSMs, B) peaks lost in mediocre single cell PSMs.

Although the isotopic envelope is more commonly used in MS1 quantitation algorithms, some MS2 peptide identification algorithms consider the +1 isotopic peak in their scoring^15^. Given the loss of mono-isotopic peaks noted above, we investigated how often an annotated *y* ion had a +1 isotopic peak. If these peaks also decline in number, the scoring functions would yield a poorer score and the PSM probability would migrate towards non-significance. We calculated how many PSMs had at least 50% of all *y* ions observed with a +1 isotopic peak. In bulk data, this is 66%. In 2ng data, this was 51% for quality PSMs compared to only 32% for mediocre PSMs. In 0.2ng data it was 39% for quality PSMs and 25% for mediocre PSMs. We also note the significant number of spectra for which there were almost no +1 isotopes (20% or fewer of the *y* ions). In 2ng quality PSMs, this was 15% compared to 41% for mediocre PSMs. In 0.2ng data it was 21% in quality PSMs and 44% in mediocre PSMs.

Both the loss of y-ions peaks and their associated ^13^C isotopic peaks are expected considering the overall loss of peaks in a spectrum. In a typical spectrum from bulk data, the average number of peaks is 682. This number is significantly less for single cell data: 104 peaks in 2ng spectra, and 86 in 0.2ng spectra.

### Intensity compression

A quality PSM is typically expected to annotate intense peaks in the spectrum, as opposed to peaks at a noise/background intensity. This idea is the basis for spectrum pre-processing filters (e.g., keep top10 peaks per 100 m/z) as well as some machine learning rescoring metrics (e.g., % intensity explained). Because single cell experiments have dramatically fewer ions, we hypothesized that the intensity of fragment ion peaks would be lower and close to non-annotated peaks. We investigated this potential intensity compression by examining the ratio between the top3 most intense annotated peaks and other peaks in the spectrum (see Figure 1). The top3/bottom3 metric, or T/B, compares the top3 and bottom3 annotated peaks. The top3/median metric, or T/M, compares the top3 annotated peaks to the median non-annotated peak.

To calculate how much of the intensity range is compressed, we compare T/B ratios for PSMs of the same peptide from bulk and single cell data. Referring back to Figure 1 as an example, the bulk spectrum has T/B = 27 and the quality 2ng spectrum has T/B = 5. Thus the intensity compression for this PSM is calculated as 27/5 = 5.4. When considering the mediocre 2ng PSM, the intensity compression is 27/2 = 13.5. A high intensity compression metric indicates that the PSM has lost a large amount of intensity range, or that the intensities of peaks are compressed. A low score for this metric indicates that the intensity range of the single cell PSM is similar to that of the PSM from bulk proteomics data.

We calculate the intensity compression metric for all peptides (Figure 3A). The comparison of bulk to quality single cell data centers around 2, which means that the intensity range in bulk data is twice as large as the intensity range in single cell data. This is true for PSMs from both 2ng and 0.2ng data. However, the comparison of bulk to mediocre single cell data centers around 4, and has a wider tail than the quality data. Therefore, the intensity range for PSMs with a mediocre q-value is far more compressed than in quality single cell PSMs.

**Figure 3.**
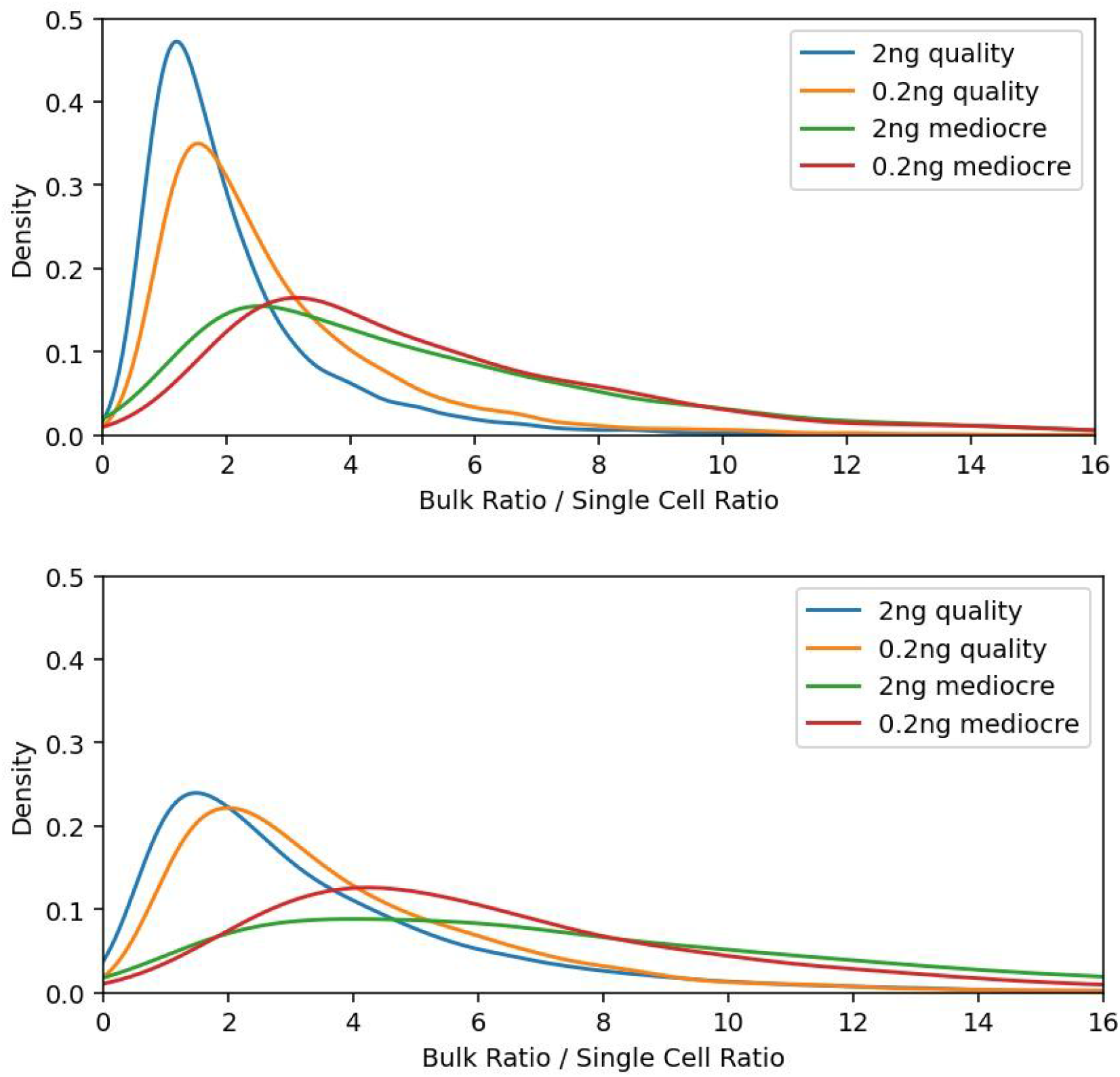
Intensity compression in single cell data. Plots show loss of intensity range in a PSM, called intensity compression. X axis is the ratio of bulk intensity range and single cell intensity range. A high intensity compression metric indicates that the PSM has lost a large amount of intensity range. The top panel measures intensity compression by comparing the top3 annotated peaks to the bottom3 annotated peaks. The bottom panel measures intensity compression by comparing the top3 annotated peaks to the median of all nonannotated peaks, a proxy for background intensity.

In addition to examining the intensity compression as measured by the top3/bottom3 ratio (T/B), we also were interested in whether the annotated peaks began to have similar intensity as non-annotated background peaks. In a typical bulk spectrum, there are hundreds of peaks and the vast majority are non-annotated and have very low intensity. These peaks are often treated as noise and may be filtered out prior to spectrum identification either by a file conversion tool like msconvert, or directly as part of a peptide identification tool. However, in single cell spectra, we noticed that the intensity of annotated *y* ion peaks begins to approach these noise peaks, as calculated by intensity compression metric using the T/M ratio (Figure 3B).

### Spectral Consistency

As seen in Figure 1, spectra from single cell datasets are distinct from traditional bulk proteomics. Single cell spectra lose peaks and signal intensity, and in mediocre spectra these losses are amplified. Given the distinction between single cell spectra and bulk spectra, another important fundamental assumption for many algorithms that needs to be evaluated is whether the fragmentation pattern remains consistent. Moreover, we were uncertain whether these low ion counts would impact the similarity of single cell spectra compared to a bulk PSM for the same peptide. We calculated the cosine similarity of all PSMs for the same peptide(see Methods). Figure 4 shows the distribution of cosine scores for each comparison, broken down into several categories. First, comparing quality single cell spectra to each other yields high similarity scores (shown by the blue, orange, and green lines), which shows that quality single cell spectra are internally consistent. We also see high scores when comparing quality spectra within only bulk data (red line). Overall this tells us that spectra of the same datatype are similar to each other. As expected, low scores are seen when comparing 2ng quality spectra to 2ng mediocre spectra (brown line). Interestingly, we also see low scores when comparing 2ng quality spectra to bulk quality spectra. The distribution (purple line) is weighted towards 0, indicating that single cell spectra and bulk spectra are dissimilar.

**Figure 4.**
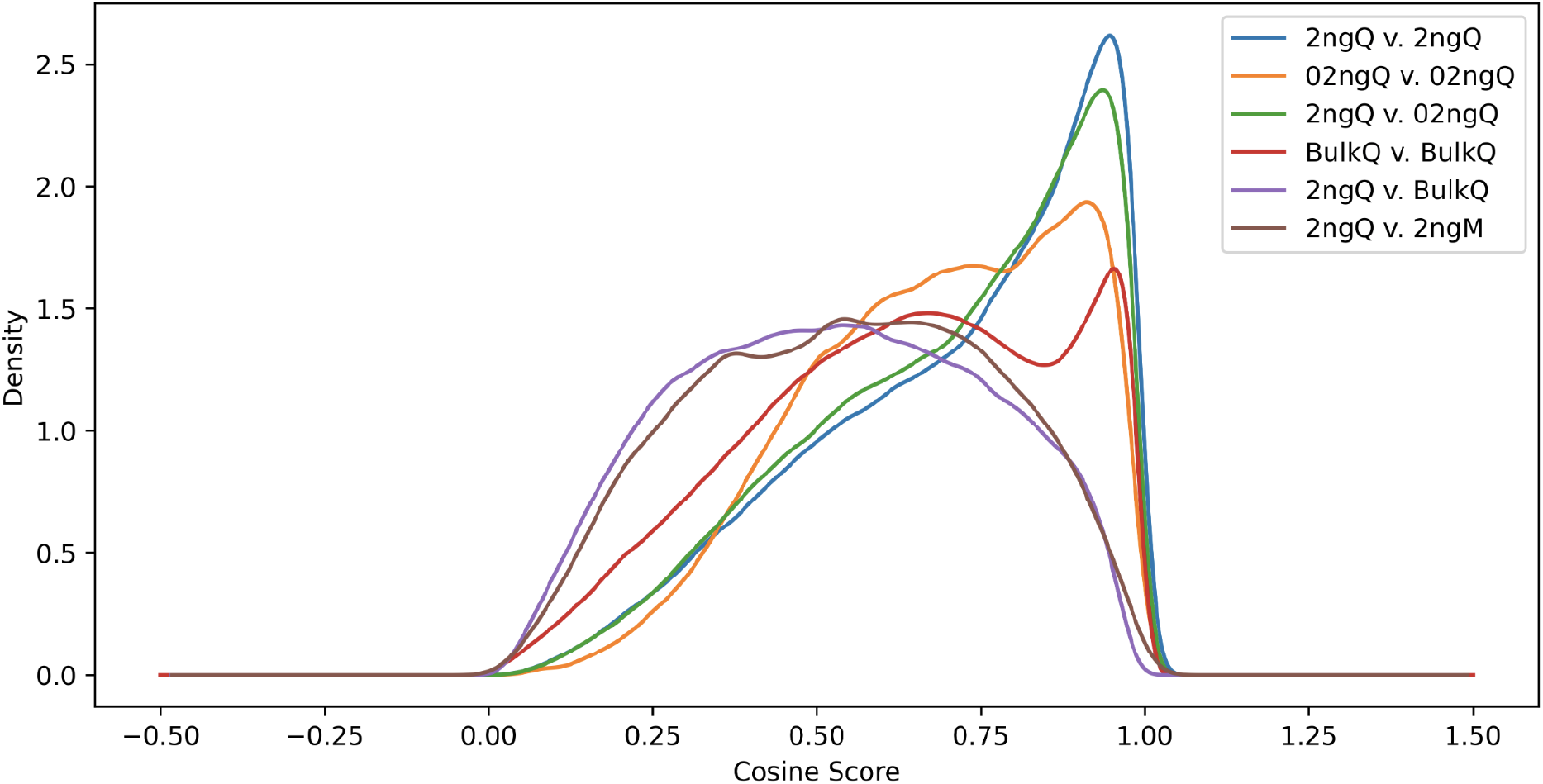
Spectral similarity for PSMs of the same peptide. The cosine similarity of PSM pairs was computed and the histogram shows the distribution of these scores. Results are grouped into six types of comparisons: 2ng quality versus 2ng quality (blue), 0.2ng quality versus 0.2ng quality (orange), bulk quality versus bulk quality (red), 2ng quality versus 2ng mediocre (brown), 2ng quality versus 0.2ng quality (green), and 2ng quality versus bulk quality (purple). Comparisons where both PSMs are from the same data type tend to have high similarity. However, comparing 2ng quality data to either 2ng mediocre or bulk quality resulted in low similarity scores.

## Discussion

Algorithms for peptide identification rely on predictable features of fragment spectra^12^. Researchers in the emerging field of single cell proteomics have been interpreting their spectra using existing tools and algorithms that were developed for bulk data, on the assumption that features remain the same for single cell spectra. We tested this assumption by analyzing three features that algorithms commonly rely on and have shown that they are different in spectra produced from single cell data.

Having a sufficient number of annotated peaks is crucial to all peptide identification algorithms, and we have shown that single cell spectra have fewer than bulk spectra. In addition to fewer annotated peaks, these spectra also have fewer total peaks, which implicitly affects the scoring of PSMs in many algorithms. For example, both XCorr and ΔC_n_ from SEQUEST implicitly rely on a high density of peaks in a spectrum. Similarly, MSGF and X!Tandem both calculate a probability score that compares the score of the top candidate peptide to the distribution of other peptides. A spectrum with significantly fewer total ions will have an altered score distribution of non-winning candidate peptides.

Beyond the simple presence of peaks, the shape of fragmentation spectra is an important feature and frequently used in algorithms for peptide identification. We show that the majority of spectra from single cell data have significant intensity compression, blurring the line between signal and noise. Additionally, spectral library searches rely on peptides producing fragment spectra which are consistent and recent work with deep learning attempts to predict the fragmentation pattern for use in a library like search^19^. We found that although spectra from single cell proteomics are internally consistent, they are distinct from spectra from bulk proteomics. This suggests that single cell spectrum identification won’t perform as well with bulk libraries and prediction methods, and that a possible improvement is to use spectral libraries built from single cell samples, and train spectrum prediction algorithms on single cell data. Understanding and accounting for the differences between bulk and single cell spectra is an important step in optimizing algorithms for single cell proteomics.

## Acknowledgements

The authors wish to thank Mingxun Wang (UCSD) for helpful discussions. This work was funded through a Sponsored Research Agreement from Biogen, Inc.

## Methods

### Data Sets

Three replicates of bulk data were obtained from publicly available analyses of HeLa digest (PRIDE project PXD011163). Single cell data were obtained from HeLa protein digest standard as described below, and are available on the ProteomeXchange partner repository MassIVE (MSV000087689). The single cell data set contains 6 runs of 2 ng data (equivalent to 10 cells), and 6 runs of .02 ng data (equivalent to 1 cell).

### Reagents and Materials

Pierce™ HeLa protein digest standard and formic acid were purchased from Thermo Fisher Scientific (Waltham, MA). Mobile phase A (0.1% formic acid in water) and mobile phase B (0.1% formic acid in acetonitrile) were respectively prepared from LC-MS grade water and acetonitrile purchased from Honeywell (Charlotte, NC). The digest standard was reconstituted to a final concentration of 200 ng/µL with 100 µL of mobile phase A to form a stock solution. For the experiments, the stock samples were further diluted to 0.2 and 2 ng/µL using the same mobile phase.

Columns: 30-µm-i.d. fused silica capillary columns from Polymicro (Phoenix, AZ) were packed with different materials: Jupiter C18 3.0 µm, 300 Å particles and Kinetex C18 core shell particles of 1.3 µm, 100 Å µm were purchased from Phenomenex (Torrance, CA); BEH C18, 1.7 µm, 130 Å was from Waters (Milford, MA). Column lengths were adjusted to keep the pressure and the linear velocity constant for all columns. The lengths were 50, 9 and 16 cm for Jupiter, Kinetex and BEH columns respectively.

Solid-phase-extraction (SPE) columns were prepared by packing Jupiter C18 particles into 100-µm-i.d. × 5-cm-long fused silica capillaries.

### LC

Previously reported by Liang et al^20^, the setup consisted of a set of Ultimate 3000 series nanoPump (NCP-3200RS) and autosampler (WPS-3000 TPL) from Thermo Fisher (Waltham, MA), a 10-port nanovolume valve (part #C72MX-4570, VICI, Houston, TX), a SPE and home-packed columns. For the autosampler, instead of using the µL pick-up, in which the sample volume is less than the total volume of the sample loop, and the rest of the sample loop is filled with mobile phase A described in a previous study^20^, we used the full loop method. Hela protein digest standard was diluted into 0.2 and 2 ng/µL range, and the sample loop size was exactly 1 µL. The sample loop was overfilled with 2.4 µL before the valve was switched to load the sample onto the SPE. The sample was loaded onto the SPE at 0.4 µL/min for 4 min at 2% B before the valve was switched again to connect the SPE and the analytical column and start the separation gradient. Flow rate was about 40 nL/m through the analytical column. The LC was coupled to an Orbitrap Exploris 480 (Thermo Fisher, Waltham, MA) as the end detector.

### Separation gradient

After the valve was switched to deliver the sample onto the analytical column, mobile phase B was increased from 2% to 8% in 3 min. A 60 min linear gradient of 8 to 25% B was used for separation, followed by a 20 min linear gradient of 22−45% B, and 45 to 90% B in 5 min. The gradients were capped at 90% B for 5 min before performing a saw tooth pattern to flush the column (switching back and forth between 2% B and 90% B in short periods). The column was re-equilibrated for 25 min at 98% mobile phase A.

### Mass Spectrometry

A Nanospray Flex ion source was used for the Orbitrap Exploris 480. The electrospray voltage was set at 2.2 kV and the temperature of the ion transfer tube was set to 200 °C. The RF lens was set at 50%.

For MS1, the Orbitrap resolution was set to 120,000 (at m/z 200) with the normalized AGC target of 300%. The scan range was between 375-1575 m/z. Maximum injection time was set at 118 ms for both 2 and 0.2 ng Hela protein digest standard. Additional filters included an intensity threshold of 5.0 e3; charge state of 2 to 5; dynamic exclusion of 30 s with mass tolerance at 10 ppm. Cycle time was set to 1.5 s.

For MS2, the isolation window was at 1.6 m/z. HCD collision energy was at 30%. The resolution was set to 30,000 and 60,000 for 2 and 0.2 ng Hela protein digest standard respectively. AGC target was 200% for both experiments. Maximum injection time was set at 118 ms for 2 ng and 246 ms for 0.2 ng sample.

### Peptide Identification

All software code and essential files for the analysis within this manuscript are found in our public GitHub repository: https://github.com/PayneLab/SingleCellSpectra.

MetaMorpheu^21^ was used for peptide identification, and the exact parameters used can be found in our GitHub repository (∼/data). Briefly, we used default parameters except for the following: number of modifications per peptide = 2, length of peptide = 8-30, Tolerance = 20 ppm, fixed modifications = carbamidomethyl, variable modifications = oxidation. The human proteome was downloaded from UniProt. The fasta file used by MetaMorpheus is uploaded to our GitHub repository in the ∼/data directory. Three tasks were used: a search task, a calibration task, and final search task. Peptide/spectrum matches used in this analysis were found in the MetaMorpheus .psmtsv files in the output folder Individual File Results. These are stored in our GitHub repository and are streamed in our analysis via python loading scripts (see ∼/data_loader.py). ‘QValue’ was used to determine the cutoff of quality data (qvalue < 1%) and mediocre data (between 1% and 20%).

Throughout the manuscript when counting PSMs, both preloading and washing/reconditioning portions of the chromatography gradient were removed based on manual inspection of acquisition rate, so that only PSMs acquired from the analytical portion of the chromatography gradient were kept. The LC-MS runtime for each file is approximately 140 minutes total, with an 80 minute analytical gradient (Supplemental Figure 2). For 2ng datasets, we begin counting PSMs when there are >= 300 scans collected per minute and we stop when there are <200 scans per minute. For the 0.2ng files, we begin when there are >= 200 scans per minute and stop when there are <100 scans per minute. No filter was applied to bulk data. The specific filter applied to each file can be found in the ∼/make_supplemental_figure2--psm_over_time.ipynb notebook.

### Spectral Features

All of the software used in the calculation and analysis of spectral features are in publicly available code within our GitHub repository https://github.com/PayneLab/SingleCellSpectra. All code necessary for the creation of each individual figure is also in the repository.

#### Calculating peaks loss

Using the MetaMorpheus PSM output file, we parse directly the annotated *y* peaks for each PSM. For each peptide, the rounded mean number of annotated *y* peaks across all PSMs is reported. The number of missing peaks is calculated by comparing the number of peaks from bulk spectra to either 2ng or 0.2ng spectra. For calculating +1 isotopic peaks, 1.0034 was added to the monoisotopic peak, with a tolerance of .01.

#### Intensity Compression

There are two metrics used to describe intensity compression: T/B and T/M. T is the mean abundance of the top 3 annotated peaks in a spectrum. B is the mean abundance of the bottom 3 annotated peaks in a spectrum. M refers to the median abundance of all non-annotated peaks in a spectrum. T/B and T/M are simple ratios averaged across all scans that identify the same peptide. In Figure 3, we plot the change in T/B from bulk data to 2ng or 0.2ng data. This is simply a ratio of ratios, T/B_bulk_ / T/B_single_cell_ .

#### Spectral Consistency

Cosine similarity scores were calculated using the calc_cosine_score.R script from the OrgMassSpecR library. The code for calculating similarity can be found in our GitHub repository in the ∼/make_figure4--spectral_consistency.ipynb.

## Supplemental Figures

**Supplemental Figure 1.**
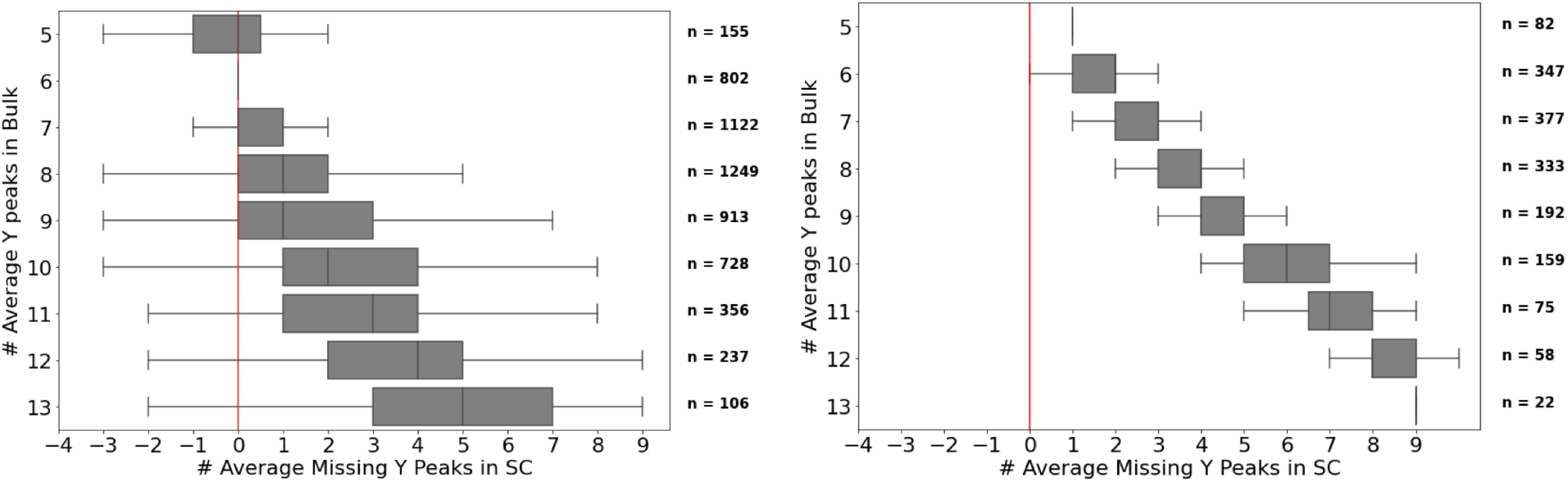
*Y* ion loss in 0.2 ng single cell spectra. This was calculated exactly like Figure 1, but with 0.2 ng data rather than 2 ng data. The results are just as what was seen in Figure 1; in quality data we lose y ions in larger peptides, and in mediocre spectra we consistently lose half of all y peaks compared to bulk spectra.

**Supplemental Figure 2.**
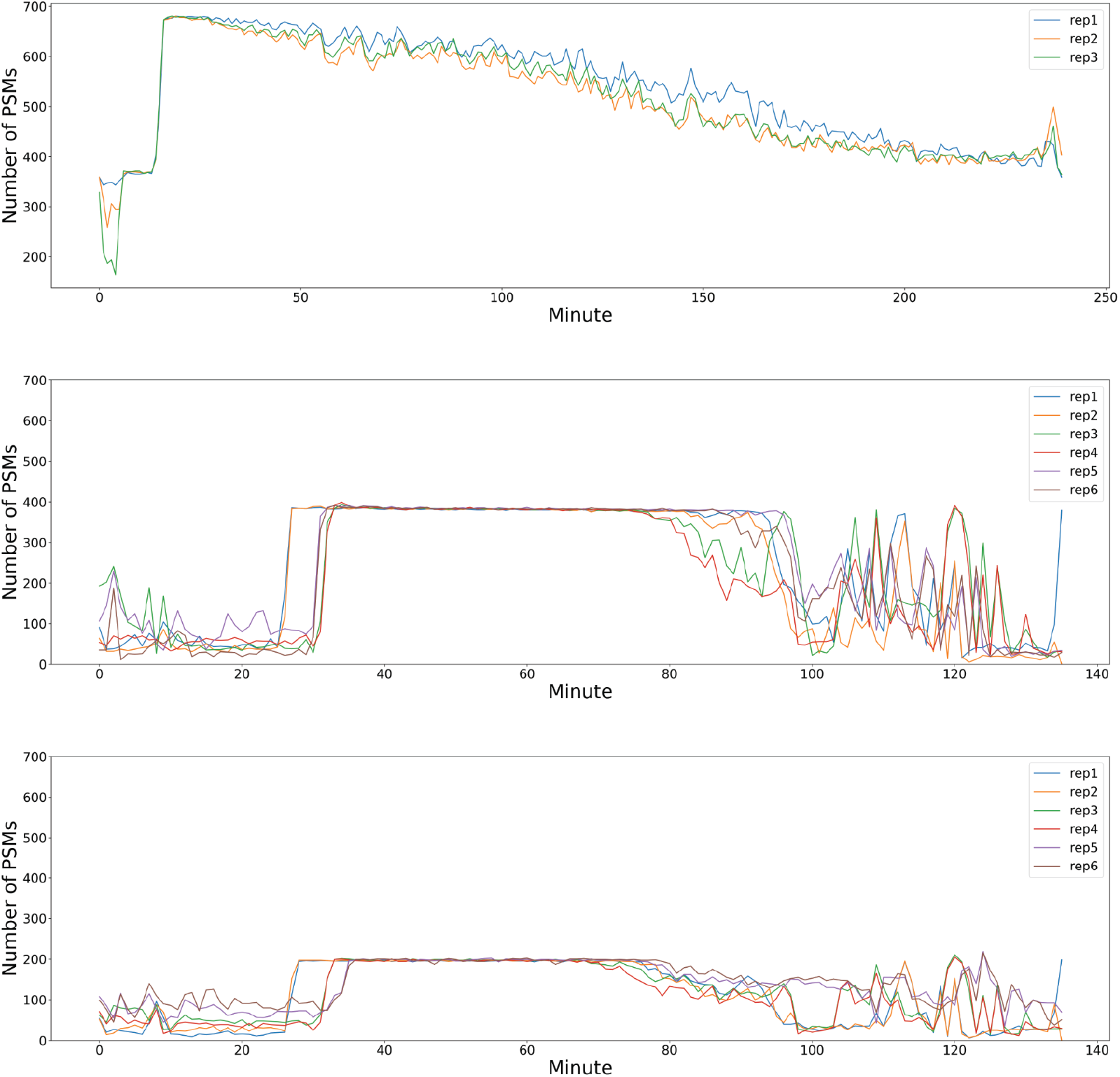
Number of PSMs acquired across the LC gradient. Each panel shows how many PSMs were collected per minute across the LC gradient. The top panel shows the gradient for each replicate of bulk data, the middle panel shows the gradient for 6 replicates of 2ng data, and the bottom panel shows 6 replicates of 0.2ng data. In single cell data the analytical part of the gradient starts ∼30 minutes in and ends after ∼90 minutes.

## References

(1) Kelly, R. T. Single-Cell Proteomics: Progress and Prospects. Mol. Cell. Proteomics MCP 2020, 19 (11), 1739–1748. https://doi.org/10.1074/mcp.R120.002234.

(2) Ctortecka, C.; Mechtler, K. The Rise of Single-cell Proteomics. Anal. Sci. Adv. 2021, 2 (3–4), 84–94. https://doi.org/10.1002/ansa.202000152.

(3) Schoof, E. M.; Furtwängler, B.; Üresin, N.; Rapin, N.; Savickas, S.; Gentil, C.; Lechman, E.; auf dem Keller, U.; Dick, J. E.; Porse, B. T. Quantitative Single-Cell Proteomics as a Tool to Characterize Cellular Hierarchies; preprint; Systems Biology, 2019. https://doi.org/10.1101/745679.

(4) Cong, Y.; Motamedchaboki, K.; Misal, S. A.; Liang, Y.; Guise, A. J.; Truong, T.; Huguet, R.; Plowey, E. D.; Zhu, Y.; Lopez-Ferrer, D.; Kelly, R. T. Ultrasensitive Single-Cell Proteomics Workflow Identifies >1000 Protein Groups per Mammalian Cell. Chem. Sci. 2021, 12 (3), 1001–1006. https://doi.org/10.1039/D0SC03636F.

(5) Dou, M.; Clair, G.; Tsai, C.-F.; Xu, K.; Chrisler, W. B.; Sontag, R. L.; Zhao, R.; Moore, R. J.; Liu, T.; Pasa-Tolic, L.; Smith, R. D.; Shi, T.; Adkins, J. N.; Qian, W.-J.; Kelly, R. T.; Ansong, C.; Zhu, Y. High-Throughput Single Cell Proteomics Enabled by Multiplex Isobaric Labeling in a Nanodroplet Sample Preparation Platform. Anal. Chem. 2019, 91 (20), 13119–13127. https://doi.org/10.1021/acs.analchem.9b03349.

(6) Virant-Klun, I.; Leicht, S.; Hughes, C.; Krijgsveld, J. Identification of Maturation-Specific Proteins by Single-Cell Proteomics of Human Oocytes. Mol. Cell. Proteomics 2016, 15 (8), 2616–2627. https://doi.org/10.1074/mcp.M115.056887.

(7) Williams, S. M.; Liyu, A. V.; Tsai, C.-F.; Moore, R. J.; Orton, D. J.; Chrisler, W. B.; Gaffrey, M. J.; Liu, T.; Smith, R. D.; Kelly, R. T.; Pasa-Tolic, L.; Zhu, Y. Automated Coupling of Nanodroplet Sample Preparation with Liquid Chromatography-Mass Spectrometry for High-Throughput Single-Cell Proteomics. Anal. Chem. 2020, 92 (15), 10588–10596. https://doi.org/10.1021/acs.analchem.0c01551.

(8) Tsai, C.-F.; Zhao, R.; Williams, S. M.; Moore, R. J.; Schultz, K.; Chrisler, W. B.; Pasa-Tolic, L.; Rodland, K. D.; Smith, R. D.; Shi, T.; Zhu, Y.; Liu, T. An Improved Boosting to Amplify Signal with Isobaric Labeling (IBASIL) Strategy for Precise Quantitative Single-Cell Proteomics. Mol. Cell. Proteomics MCP 2020, 19 (5), 828–838. https://doi.org/10.1074/mcp.RA119.001857.

(9) Cheung, T. K.; Lee, C.-Y.; Bayer, F. P.; McCoy, A.; Kuster, B.; Rose, C. M. Defining the Carrier Proteome Limit for Single-Cell Proteomics. Nat. Methods 2021, 18 (1), 76–83. https://doi.org/10.1038/s41592-020-01002-5.

(10) Sun, B.; Kovatch, J. R.; Badiong, A.; Merbouh, N. Optimization and Modeling of Quadrupole Orbitrap Parameters for Sensitive Analysis toward Single-Cell Proteomics. J. Proteome Res. 2017, 16 (10), 3711–3721. https://doi.org/10.1021/acs.jproteome.7b00416.

(11) Eng, J. K.; McCormack, A. L.; Yates, J. R. An Approach to Correlate Tandem Mass Spectral Data of Peptides with Amino Acid Sequences in a Protein Database. J. Am. Soc. Mass Spectrom. 1994, 5 (11), 976–989. https://doi.org/10.1016/1044-0305(94)80016-2.

(12) Dancík, V.; Addona, T. A.; Clauser, K. R.; Vath, J. E.; Pevzner, P. A. De Novo Peptide Sequencing via Tandem Mass Spectrometry. J. Comput. Biol. J. Comput. Mol. Cell Biol. 1999, 6 (3–4), 327–342. https://doi.org/10.1089/106652799318300.

(13) Ma, B.; Zhang, K.; Hendrie, C.; Liang, C.; Li, M.; Doherty-Kirby, A.; Lajoie, G. PEAKS: Powerful Software for Peptide de Novo Sequencing by Tandem Mass Spectrometry. Rapid Commun. Mass Spectrom. RCM 2003, 17 (20), 2337–2342. https://doi.org/10.1002/rcm.1196.

(14) Frank, A.; Pevzner, P. PepNovo: De Novo Peptide Sequencing via Probabilistic Network Modeling. Anal. Chem. 2005, 77 (4), 964–973. https://doi.org/10.1021/ac048788h.

(15) Payne, S. H.; Yau, M.; Smolka, M. B.; Tanner, S.; Zhou, H.; Bafna, V. Phosphorylation-Specific MS/MS Scoring for Rapid and Accurate Phosphoproteome Analysis. J. Proteome Res. 2008, 7 (8), 3373–3381. https://doi.org/10.1021/pr800129m.

(16) Käll, L.; Canterbury, J. D.; Weston, J.; Noble, W. S.; MacCoss, M. J. Semi-Supervised Learning for Peptide Identification from Shotgun Proteomics Datasets. Nat. Methods 2007, 4 (11), 923–925. https://doi.org/10.1038/nmeth1113.

(17) Kim, S.; Pevzner, P. A. MS-GF+ Makes Progress towards a Universal Database Search Tool for Proteomics. Nat. Commun. 2014, 5, 5277. https://doi.org/10.1038/ncomms6277.

(18) Giansanti, P.; Strating, J. R. P. M.; Defourny, K. A. Y.; Cesonyte, I.; Bottino, A. M. S.; Post, H.; Viktorova, E. G.; Ho, V. Q. T.; Langereis, M. A.; Belov, G. A.; Nolte-’t Hoen, E. N. M.; Heck, A. J. R.; van Kuppeveld, F. J. M. Dynamic Remodelling of the Human Host Cell Proteome and Phosphoproteome upon Enterovirus Infection. Nat. Commun. 2020, 11 (1), 4332. https://doi.org/10.1038/s41467-020-18168-3.

(19) Tiwary, S.; Levy, R.; Gutenbrunner, P.; Salinas Soto, F.; Palaniappan, K. K.; Deming, L.; Berndl, M.; Brant, A.; Cimermancic, P.; Cox, J. High-Quality MS/MS Spectrum Prediction for Data-Dependent and Data-Independent Acquisition Data Analysis. Nat. Methods 2019, 16 (6), 519–525. https://doi.org/10.1038/s41592-019-0427-6.

(20) Liang, Y.; Acor, H.; McCown, M. A.; Nwosu, A. J.; Boekweg, H.; Axtell, N. B.; Truong, T.; Cong, Y.; Payne, S. H.; Kelly, R. T. Fully Automated Sample Processing and Analysis Workflow for Low-Input Proteome Profiling. Anal. Chem. 2021, 93 (3), 1658–1666. https://doi.org/10.1021/acs.analchem.0c04240.

(21) Solntsev, S. K.; Shortreed, M. R.; Frey, B. L.; Smith, L. M. Enhanced Global Post-Translational Modification Discovery with MetaMorpheus. J. Proteome Res. 2018, 17 (5), 1844–1851. https://doi.org/10.1021/acs.jproteome.7b00873.

